# Phospholipid-Cellulose Interactions: Insight from Atomistic Computer Simulations for Understanding the Impact of Cellulose-Based Materials on Plasma Membranes

**DOI:** 10.1101/425686

**Authors:** Andrey A. Gurtovenko, Evgenii I. Mukhamadiarov, Andrei Yu. Kostritskii, Mikko Karttunen

## Abstract

Cellulose is an important biocompatible and nontoxic polymer widely used in numerous biomedical applications. The impact of cellulose-based materials on cells and, more specifically, on plasma membranes that surround cells, however, remains poorly understood. To this end, here we performed atomic-scale molecular dynamics (MD) simulations of phosphatidylcholine (PC) and phosphatidylethanolamine (PE) bilayers interacting with the surface of a cellulose crystal. Both biased umbrella sampling and unbiased simulations clearly show the existence of strong attractive interactions between phospholipids and cellulose: the free energy of the cellulose-bilayer binding was found to be −1.89 and −1.96 kJ/mol per cellulose dimer for PC and PE bilayers, respectively. Although the values are similar, there is a pronounced difference between PC and PE bilayers. The driving force in both cases is the formation of hydrogen bonds. There are two distinct types of hydrogen bonds: 1) between the lipid head groups and the hydroxyl (hydroxymethyl) groups of cellulose, and 2) lipid-water and cellulose-water bonds. The former is the dominant component for PE systems whereas the latter dominates in PC systems. This suggests that achieving controlled binding via new cellulose modifications must pay close attention to the lipid head groups involved. The observed attractive phospholipid-cellulose interactions have a significant impact on bilayer properties: a cellulose crystal induces noticeable structural perturbations on the bilayer leaflet next to the crystal. Given that such perturbations can be undesirable when it comes to the interactions of cellulose-based materials with cell membranes, our computational studies suggest that the impact of cellulose could be reduced through chemical modification of the cellulose surface which prevents cellulose-phospholipid hydrogen bonding.

## INTRODUCTION

Cellulose is a biodegradable and easily accessible natural polymer that is widely used in industry and diverse applications.^1–5^ Furthermore, cellulose is both biocompatible and nontoxic, which additionally promotes its applications in medicine, for example in wound dressings, bone implants and bionanocomposites with antimicrobial activity.^6–10^ Despite such a wide range of applications, the impact and interactions of cellulose with cells remain to be fully understood. Here, we focus on the interactions of cellulose with model cellular plasma membranes.

We employ atomic-scale MD simulations to obtain a detailed insight into the structure and properties of the interfacial region between the surface of a cellulose nanocrystal and model phospholipid membranes. We chose to consider phosphatidylcholine (PC) and phosphatidylethanolamine (PE) membranes, since PC and PE lipids are the main representatives of zwitterionic lipids in eukaryotic plasma membranes.^11-13^ Our preliminary studies showed^14^ that one could anticipate strong interactions between the cellulose crystal and the PC lipid bilayer. However, no definitive conclusions could be drawn on the basis of the model employed in Ref. 14 as it suffered from limitations inherent for lipid bilayers on a solid support.^15–17^ In particular, a supported bilayer could not reach the equilibrium bilayer-support distance due the fact that bulk water did not have access to the region between the bilayer and the solid (cellulose) support.^14^

To overcome these limitations, here we have improved our model by considering a cellulose crystal of a finite size, whose surface area was set to be smaller than the area of the lipid bilayer, Figure 1. In this case water molecules are free to move in and out of the interfacial cellulose-bilayer region. In a way, our model is similar to the so-called semi-supported bilayers in which a lipid bilayer is placed on top of a nanoscopically-structured (porated) support.^18,19^

**Figure 1:**
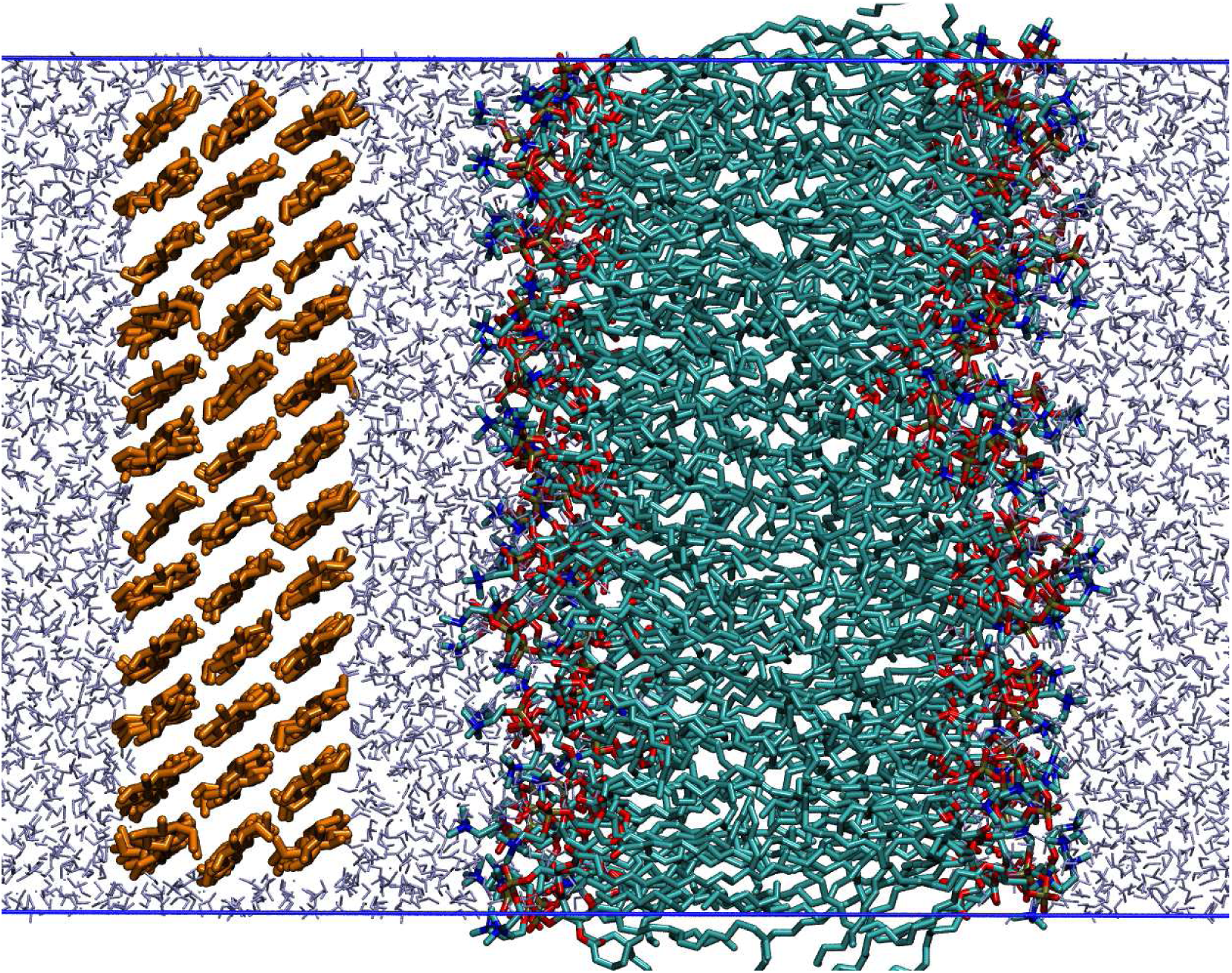
A snapshot of a cellulose-bilayer system considered in this study (the system POPC-CEL is shown). POPC lipids are shown in cyan, red and blue, a cellulose crystal in orange and water molecules in ice-blue.

The use of such a model makes it possible to properly equilibrate the cellulose-bilayer system and correspondingly provides an easy access to the equilibrium level of hydration of the interfacial region as well as to the equilibrium distance between the cellulose crystal and the phospholipid bilayer. Both these characteristics can potentially be measured experimentally. Importantly, with the use of biased umbrella sampling MD simulations we were able to evaluate for a first time the free-energy of binding of PC and PE lipid bilayers to the surface of a cellulose crystal. Our computational findings reveal strong attractive interactions between the cellulose crystal and both PC and PE bilayers due to formation of cellulose-lipid hydrogen bonds. Because of these attractive interactions, the cellulose crystal has a noticeable impact on phospholipid bilayers, resulting in an asymmetry in the properties of the opposite bilayer leaflets.

## METHODS

We have performed atomic-scale MD simulations of a finite cellulose crystal placed nearby palmitoyl-oleoyl-phosphatidylcholine (POPC) or palmitoyl-oleoyl-phosphatidyl-ethanolamine (POPE) lipid bilayers, see Figure 1.

The initial structure of the cellulose crystal was taken from our previous study.^14^ Briefly, it is based on the structure of I*β*^20^ cellulose and comprises three layers of cellulose chains (12 chains in each layer). All cellulose chains are of the same length and consist of 6 cellobiose units. Unlike in our previous paper,^14^ here the cellulose crystal has a finite size (6.63 nm × 6.61 nm in the *XY*-direction), that is, the cellulose chains are not covalently linked to their periodic images. 1n turn, a lipid bilayer has a larger size (7 nm × 7 nm) and correspondingly larger surface area as compared to the cellulose crystal (48.9 vs 43.8 nm^2^), so that water molecules can flow between cellulose crystal edges and the boundaries of a simulation box, see Figure 1. POPC and POPE lipid bilayers consisted of 148 and 168 lipids, respectively (see Table 1) due to the well-known difference^21^ in the area per lipid of POPC and POPE bilayers.

**Table 1:** Simulated cellulose-phospholipid systems.

| System | N<sub>lipids</sub> | Simulation time [ns] |
| --- | --- | --- |
| POPC | 148 | 500 |
| POPE | 168 | 500 |
| POPC-CEL | 148 | 600 |
| POPE-CEL | 168 | 600 |
| POPC-CEL-PMF | 148 | $32 \times 100$ |
| POPE-CEL-PMF | 168 | $32 \times 100$ |

In unbiased MD simulations (systems POPC-CEL and POPE-CEL in Table 1) a lipid bilayer was placed in the vicinity of a cellulose crystal in such a way that the initial distance between the centers of masses (COM) of the bilayer and the crystal was ~ 4.6-4.7 nm. The cellulose-bilayer systems were hydrated with ~ 11,500 water molecules; the total number of atoms in the systems amounted to ~ 63,800. 1n addition to the cellulose-membrane systems, we also considered free-standing lipid bilayers (POPC and POPE systems in Table 1). Such bilayers had the same number of PC and PE lipids as the cellulose-bilayer systems and were hydrated with ~5,600 water molecules; the total number of atoms was ~37,000.

The choice of force-field for a heterogeneous system (i.e. for a system containing biomolecules of different types) is not straightforward and always represents a trade-off, since it should describe in a reasonable way all the system components (lipids and carbohydrates in our case). While lipid bilayers are well described by the state-of-the-art force-fields such as CHARMM36^21^ and AMBER-like force-fields (e.g. Lipid14^22^ and Slipids^23^), the situation with polysaccharides is more involved. There is a general consensus that GLYCAM (an AMBER-like force-field for carbohydrates) and various modifications of the CHARMM force-fields have similar behavior and give similar structures.^24–26^ However, the GLYCAM force-field is not compatible with other force-fields of the AMBER family as it uses different scaling of the 1-4 interactions.^27^ Given this and also the fact that CHARMM36 is currently considered as one of the best force-fields for lipids,^28^ we chose the CHARMM force-field for simulating lipid-cellulose systems.

Therefore, the CHARMM35^29,30^ and CHARMM36^21^ force-fields were used to describe cellulose monosaccharides and phospholipids, respectively. Water was modeled by the CHARMM version of the TIP3P model. ^31^ The inner structure of the cellulose crystal was kept rigid by imposing position restraints on all heavy atoms of the monosaccharide rings with the exception of hydroxyl oxygens and exocyclic groups.

All simulations were carried out in the NPT ensemble (T = 310 K and P = 1 bar). Pressure was controlled semiisotropically and the thermostat was applied separately to cellulose, lipids and water. Each system was pre-equilibrated for 20 ns with the use of the Berendsen scheme^32^ for both thermostat and barostat. For production runs, the Nosé-Hoover thermostat^33,34^ and the Parrinello-Rahman barostat^35^ were employed. Periodic boundary conditions were applied in all three directions. All hydrogen bonds were constrained with the P-LINCS algorithm. ^36^ To handle electrostatic interactions the particle-mesh Ewald method (PME)^37^ with a real-space cutoff of 1.2 nm was used. The time step was 2fs. Unbiased MD simulations (systems POPC-CEL and POPE-CEL) were extended to 600 ns, while the simulations of free-standing lipid bilayers were 500 ns long, see Table 1. All simulations were carried out with the Gromacs 5.1.4 simulation suite.^38^

To explore the size effects, the influence of salt ions and the impact of semiisotropic pressure coupling, we performed several additional simulations for POPC-cellulose and POPE-cellulose systems. First, we considered a system with an elevated number of POPC lipids (160 lipids vs. 148 lipids presented in Table 1) and, correspondingly, with an enlarged simulation box size in the X- and Y-directions (~7.25 nm vs. ~7 nm). Second, we explored the behavior of the POPC-CEL system in the presence of NaCl salt of physiological concentration (150mM). Finally, simulations of POPC-CEL and POPE-CEL systems were repeated with the pressure coupling switched off in the bilayer plane using the NP_*z*_AT ensemble.

The umbrella sampling technique^39^ was used to evaluate the free energy of binding of a phospholipid bilayer to the surface of a cellulose crystal. First, a lipid bilayer was placed in aqueous solution parallel to the cellulose crystal at the bilayer-cellulose COM distance of 5.8 nm. The pull code supplied with the Gromacs package^38,40^ was employed to obtain starting configurations for umbrella sampling calculations. A phospholipid bilayer was slowly pulled along the reaction coordinate (the COM distance between the crystal and the bilayer in the direction perpendicular to the bilayer surface) with a velocity of 0.0001 nm/ps and a force constant of 1000 kJ mol^−1^ nm^−2^. When the bilayer established a contact with the crystal surface both the velocity and force constant were increased to 0.05 nm/ps and 3000 kJ mol^−1^ nm^−2^, respectively.

From these pulling trajectories 32 windows were extracted for umbrella sampling. The spacing between windows was 0.1 nm (from 2.7 to 5.8 nm along the reaction coordinate). Each window was simulated for 100 ns with the force constant set to 3000 kJ mol^−1^ nm^−2^. The potential of mean force (PMF) was calculated with the use of the weighted histogram analysis (WHAM)^41^ method as it is implemented in the *gmx wham* routine of the Gromacs suite.^38^ The PMF profiles converged after first 20 ns and the remaining parts of trajectories (80 ns) were used to calculate PMF. Statistical errors for PMF were estimated with the use of bootstrapping analysis.^40^

## RESULTS AND DISCUSSION

### Cellulose-Bilayer Binding and Energetics

Since the main focus of our study is on the impact of cellulose crystal on the structure and properties of lipid bilayers, one has to establish first a reference for systematic comparison. To this end, we performed simulations of free-standing lipid bilayers in aqueous solution (systems POPC and POPE in Table 1). The simulations were extended to 500 ns; last 300 ns were used for calculating the structural characteristics of the bilayers.

The area per lipid for the POPC bilayer was found to be 0.64±0.01 nm^2^ in agreement with previous simulations that employed the same lipid force-field (0.647±0.002 nm^2^, T=303K)^21^ and also with experimental data (0.66 nm^2^, T=310K).^42^ The thickness of the POPC bilayer equals 3.86±0.03 nm in line with simulations (3.88±0.01 nm, T=320K)^43^ and experimental (3.82 nm, T = 320 K)^44^ data. As for the orientation of polar head groups of POPC lipids, the angle between PN vectors and the outward bilayer normal was estimated to be 69.5±0.3 degrees (cf. 70.1 degrees reported in previous simulations).^43^

The POPE lipid bilayer is characterized by denser packing of lipids as compared to the POPC bilayer due to different chemistry of the lipid head groups.^45^ In particular, this results in smaller area per lipid and larger thickness of the bilayer. Indeed, the area per lipid for the POPE bilayer was found to be 0.55±0.01 nm^2^ (cf. 0.555±0.004 nm^2^ (T=308K) reported in simulations^43^ and 0.58 nm^2^ (T=308K) observed in experiments).^46^ The thickness of the POPE bilayer was found to be 4.28±0.05 nm (4.33±0.03 at T=308K in earlier simulations^43^ and 4.05 nm at T=308K) in experiments^46^). The PE polar head groups turned out to be more horizontally oriented than that of POPC: the PN vector equals 77.6±0.3 degrees in agreement with previous simulations that found 77.5 degrees.^43^ The structural differences between POPC and POPE bilayers can also be seen in the component-wise mass density profiles presented in Figure S1. The larger thickness of the POPE bilayer is translated to the larger distance between the peaks of the lipid’s density in the opposite leaflets, while water penetrates deeper into the POPC bilayer as compared to the POPE counterpart due the denser packing of PE lipids. Summarizing, the employed models of POPC and POPE bilayers are fully in agreement with previous independent simulation and experimental data and correctly describe the structural difference between the PC and PE lipid bilayers.

Turning now to the cellulose-bilayer systems, we focus first on the binding kinetics of a phospholipid bilayer and a cellulose crystal. To this end, a bilayer was placed nearby a crystal parallel to the crystal surface, see Figure 1. To follow the process of binding, we calculated the distance between centers of masses (COMs) of a lipid bilayer and a cellulose crystal along the *z*-axis (the bilayer normal). The corresponding curves are presented in Figure 2.

**Figure 2:**
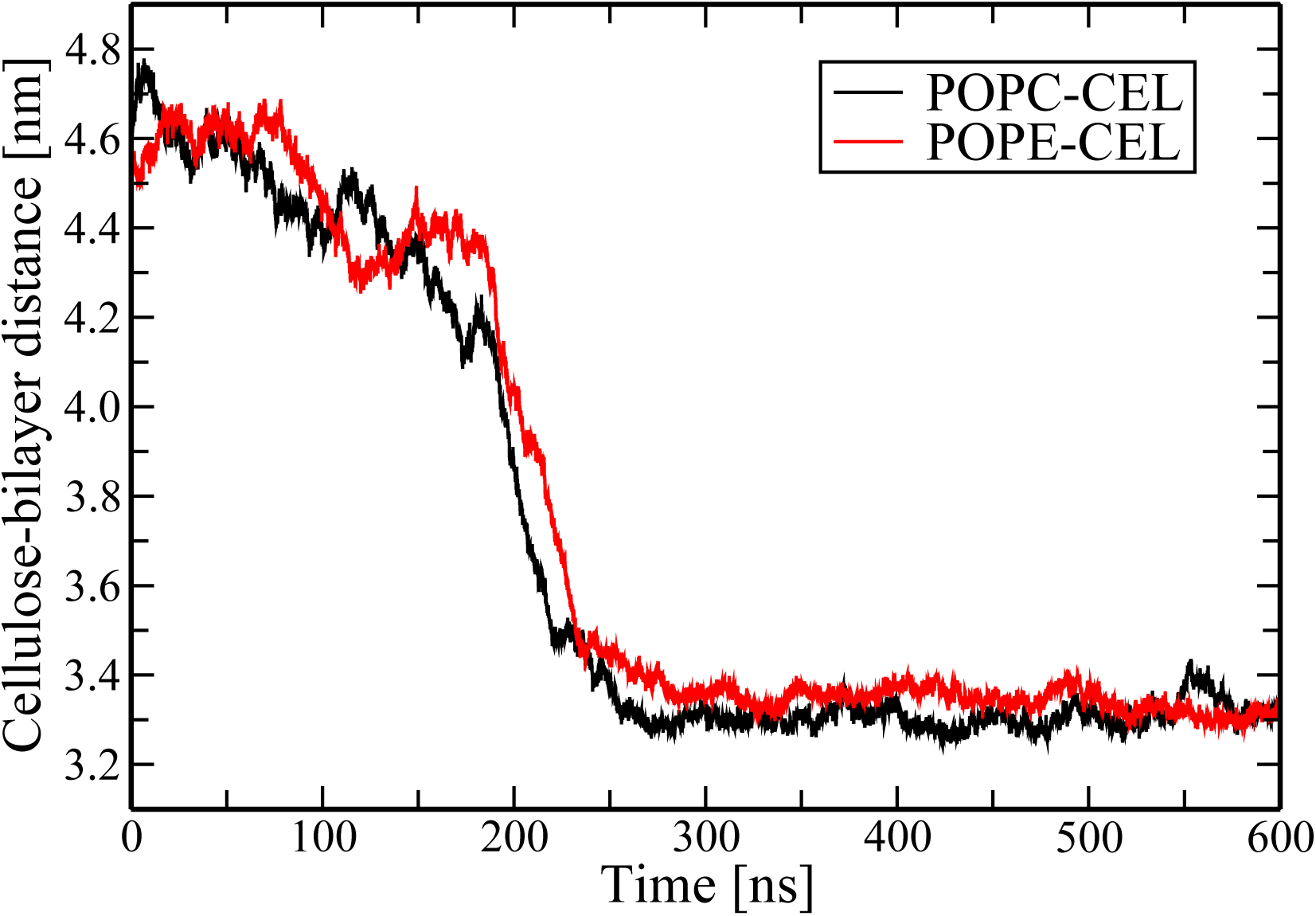
The distance between centers of mass of a phospholipid bilayer and a cellulose crystal along the *z*-axis(the bilayer normal) as a function of time. Shown are results for POPC-CEL (black line) and POPE-CEL (red line) systems.

As it is evident from Figure 2 both POPC and POPE lipid bilayers bind to the surface of a cellulose crystal. Binding occurs spontaneously: the bilayer spends around 200 ns in aqueous solution near the crystal until the first contacts with cellulose are established. After that there is a steep drop (~ 1 nm in 10-20 ns) in the COM distance between the bilayer and the crystal. Once bound to the cellulose surface, the bilayer remains in contact with the crystal for the rest of the simulation.

Interestingly, both POPC and POPE bilayers show a very similar behavior despite the different chemical structures of their polar head groups. To exclude possible artifacts related to the semiisotropic pressure coupling, we repeated unbiased simulations of POPC-CEL and POPE-CEL systems in the NP_*z*_AT ensemble. In practice, the pressure coupling was switched off in the plane of the bilayer, so that the bilayer area was kept constant during the course of the simulations. As seen in Figure S2, the semiisotropic pressure coupling has no effect on the attractive interactions between a cellulose crystal and a lipid bilayer (cf. Figure 2). For POPC bilayers we also repeated the simulations for a bilayer of a larger size (160 lipids) as well as for a POPC-cellulose system under the presence of NaCl salt of physiological concentration (150mM), see Figure S3. Overall, the same pattern of the lipid-cellulose binding is observed although salt ions slow down the binding to some extent.

Figure 2 shows that the COM bilayer-cellulose distances reach equilibrium after about 300 ns at 3.31±0.03 and 3.34±0.03 nm for POPC and POPE lipid bilayers, respectively. Here (and in the following) averaging is carried out over last 300 ns of the MD trajectories, if not stated otherwise. Note that the COM distance is not a direct measure of how close the lipid bilayer and the crystal are; the observed difference is most likely due to a smaller thickness of the POPC bilayer as compared to the POPE bilayer.^43^

Although the above findings clearly imply attractive interactions between the lipid bilayer and cellulose, unbiased simulations do not serve as a conclusive proof of attractive interactions. To investigate the interactions quantitatively, we employed umbrella sampling MD simulations to explore the energetics of the lipid-cellulose binding. Figure 3 shows the potential of mean force (PMF). The COM distance along the *z*-axis (the bilayer normal) between the lipid bilayer and the cellulose crystal was used as the reaction coordinate.

**Figure 3:**
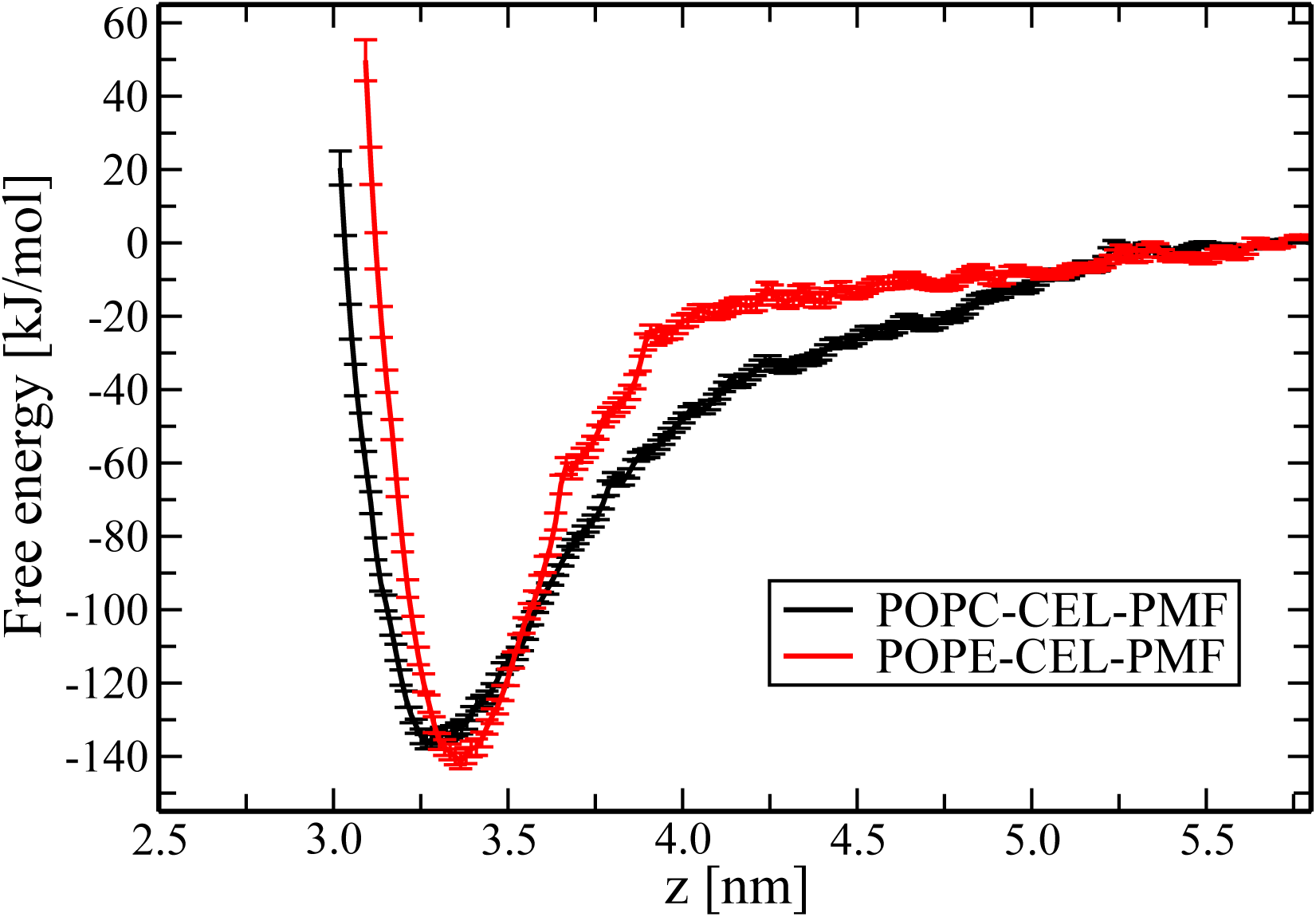
Free energy profile (potential of mean force) for binding of phospholipid bilayers from aqueous solution to the surface of a cellulose crystal. Shown are results for POPC-CEL-PMF (black line) and POPE-CEL-PMF (red line) systems. Statistical errors were estimated with the use of bootstrapping analysis.^40^

The PMF profiles clearly show that binding of POPC and POPE lipid bilayers is characterized by a deep well, that is, strong attractive interactions between phospholipids and cellulose. The depths of the free energy minima were found to be −136±2 and −141±2 kJ/mol for the POPC and POPE lipid bilayers, respectively. The PMF minima are located at 3.27 nm (the POPC-CEL-PMF system) and 3.36 nm (the POPE-CEL-PMF system), which agree well with the results of unbiased simulations, see Figure 2. The large free energy values correspond to lipid bilayer binding to a cellulose crystal patch of surface area of 43.8 nm^2^. To have an estimate for the binding energy of a single cellulose dimer, it is instructive to normalize the free energy by the number of dimers (72) on the surface of the cellulose crystal. This yields −1.89±0.03 and −1.96±0.03 kJ/mol for the free energy of binding of a cellulose dimer to POPC and POPE lipid bilayers, respectively. Thus, it can be concluded that phospholipid-cellulose binding is energetically very favorable. As we proceed to show, the origin of the observed strong binding is the formation of hydrogen bonds between the cellulose surface and the polar head groups of the lipid molecules. Furthermore, these strong attractive lipid-cellulose interactions have a noticeable impact on the properties of the lipid bilayers.

### The Structure of the Cellulose-Bilayer Interfacial Region

To explore the influence of cellulose on a lipid bilayer upon binding, we calculated the mass density profiles of the major components of the cellulose-lipid systems. The corresponding profiles are presented in Figure 4 for both POPC-CEL and POPE-CEL systems. The data clearly shows that the tight binding to cellulose has a strong impact on the structure of the lipid bilayer, resulting in a pronounced asymmetry in the density profiles of the opposite bilayer leaflets. While the distal leaflet turned out to be largely unaffected by the cellulose crystal, the proximal leaflet (i.e. leaflet next to the crystal surface) undergoes considerable structural changes. In particular, the lipid density profiles of the proximal leaflets are characterized by the appearance of two peaks in contrast to the distal leaflets where the conventional shape typical for free-standing phospholipid bilayers is observed, Figure 4.

**Figure 4:**
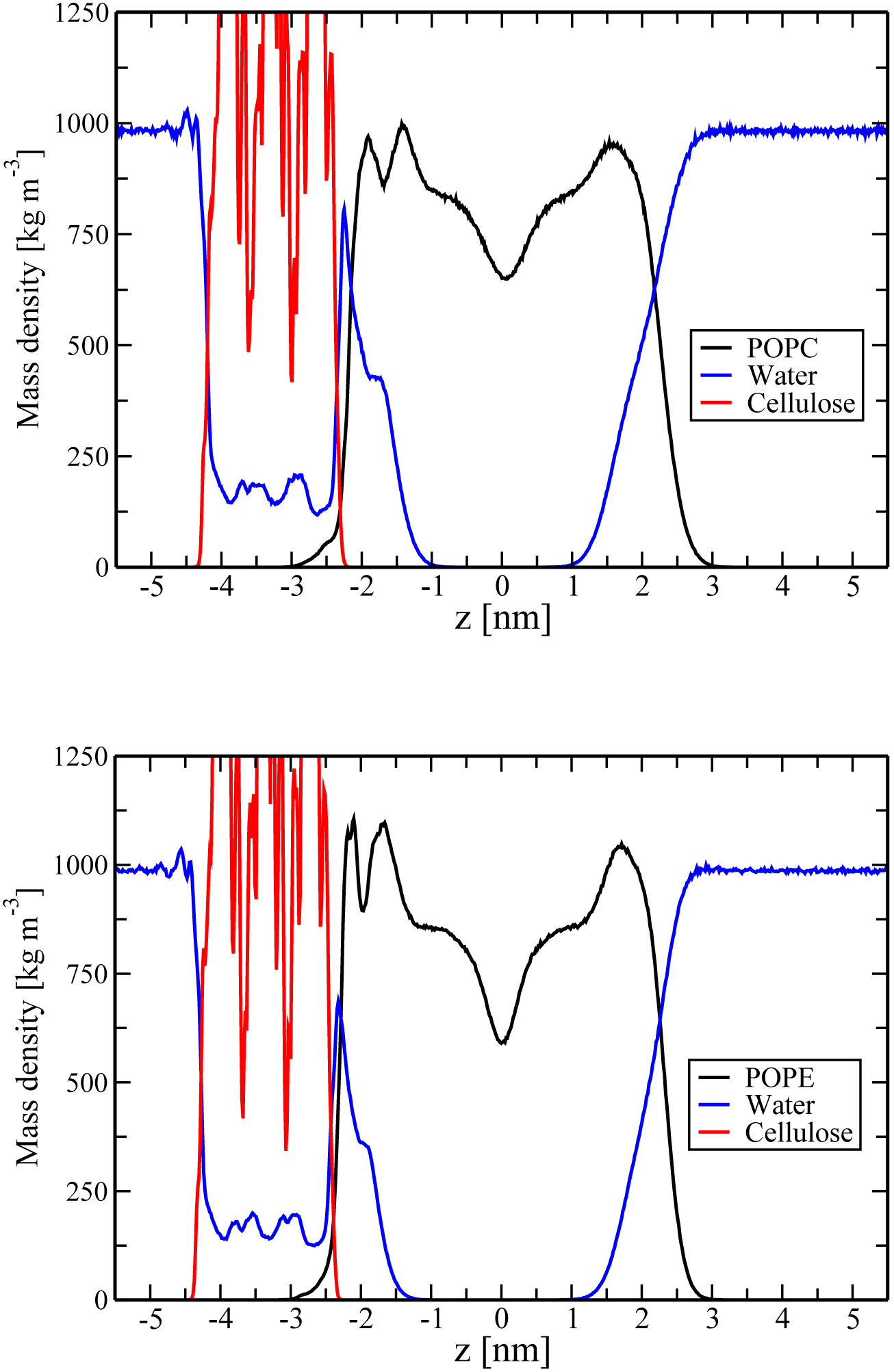
Mass density profiles for phospholipids (black line), water molecules (blue line) and cellulose (red line) for POPC-CEL (top) and POPE-CEL (bottom) systems as a function of the distance from the bilayer center (*z*=0). Note that the appearance of water molecules and lipids within the cellulose crystal density profile is due to the fact that the crystal has a finite size, so that there is free space between the crystal edges and the boundaries of a simulation box, see Figure 1.

Similar structural changes have been previously reported for POPC-cellulose systems under low hydration of the interfacial region.^14^ The results here confirm the same pattern also for POPE lipid bilayers, highlighting its universal character provided the attractive cellulose-bilayer interactions are strong enough. In more general context, several computational studies of supported lipid bilayers have witnessed similar support-induced asymmetry in the structure of the opposite lipid monolayers.^15–18^

A more detailed insight into the structure of the interfacial cellulose-lipid region can be gained via inspecting the component-wise density profiles for the principal atoms of lipid molecules and cellulose. Figure 5 shows the corresponding profiles for phosphorus, nitrogen and carbonyl oxygen atoms of the polar lipid head groups, for oxygen atoms O2(O12) and O3(O13) of the hydroxyl groups, and oxygen atoms O6(O16) of the exocyclic hydroxymethyl groups of cellulose chains, see Figure S2 for the numbering of cellulose atoms. It is noteworthy that among all key oxygen atoms of cellulose, the density peak of O6(O16) atoms is located closer to the lipid bilayer because the exocyclic hydroxymethyl groups O6(O16) (groups of carbon C6(C16)) are larger than the hydroxyl groups O2(O12) and O3(O13). Therefore, one can anticipate stronger interactions of hydroxymethyl groups O6(O16) with lipid head groups as compared to their O2(O12) and O3(O13) counterparts.

**Figure 5:**
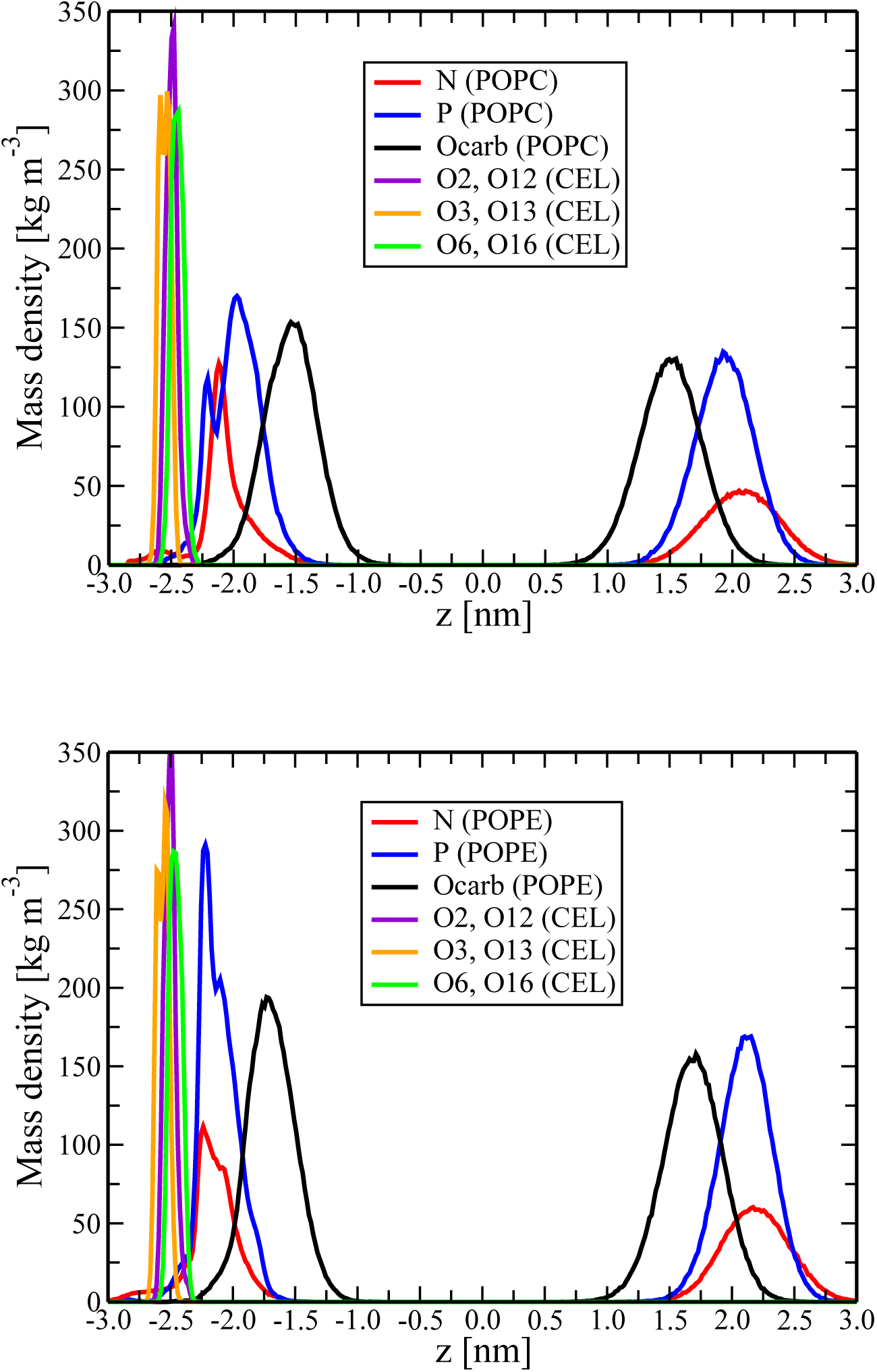
Component-wise mass density profiles for principal atoms of lipid head groups and cellulose chains as a function of the distance from the bilayer center (*z*=0). Shown are results for POPC-CEL (top) and POPE-CEL (bottom) systems.

Next, the profiles of lipid atoms of the distal and proximal leaflets are compared to shed light on how the binding of a cellulose crystal affect the fine structure of lipid head groups of POPC and POPE bilayers. In the proximal leaflets of the POPC-CEL and POPE-CEL systems the density profile peaks of the nitrogen and phosphorus atoms become narrower and higher, the effect being more pronounced for POPE bilayers. Furthermore, the peaks of the proximal leaflets are shifted toward the cellulose surface, Figure 5. All these changes in the structure of the lipid head group region are due to strong lipid-cellulose interactions. The effects of cellulose appear to be stronger in the case of the POPE lipid bilayer as compared to the POPC counterpart, leading, e.g., to a smaller equilibrium bilayer-crystal distance: 0.25 nm vs 0.47 nm for POPE-CEL and POPC-CEL systems, respectively (the distance was estimated as the distance between the main density peaks for the phosphorus atoms of the lipids and O6(O16) atoms of the hydroxymethyl groups of cellulose).

Another important structural property of a lipid bilayer is the orientation of polar head groups. Given the strong impact of cellulose on the structure of bilayer leaflets next to the crystal surface, one can anticipate an asymmetry in the orientation of lipid head groups in the opposite leaflets.^14^ Indeed, as one can see from Figure 6, the probability distributions of the angle between PN vectors of lipids and the outward bilayer normal differ considerably for proximal and distal bilayer leaflets. Overall, the distributions for proximal leaflets are shifted to larger values of the PN angle, which implies a more horizontal re-orientation of lipid head groups. Importantly, we witness the same cellulose-induced effects for both POPC and POPE lipid bilayers, see Figure 6. For POPC bilayers the average PN angle for the distal leaflet was found to be 69.3±0.1 degrees, which coincides with the value measured for a freestanding POPC bilayer, see previous Section. In turn, the binding to cellulose increases the PN angle in the proximal leaflet to 74.4±0.2 degrees. As far as the POPE-CEL system is concerned, the average PN angle was found to be 77.1±0.1 and 81.7±0.2 degrees for the distal and proximal leaflets, respectively. Again, the value of the PN angle of the distal leaflet in perfect agreement with what was reported for a free-standing POPE bilayer.

**Figure 6:**
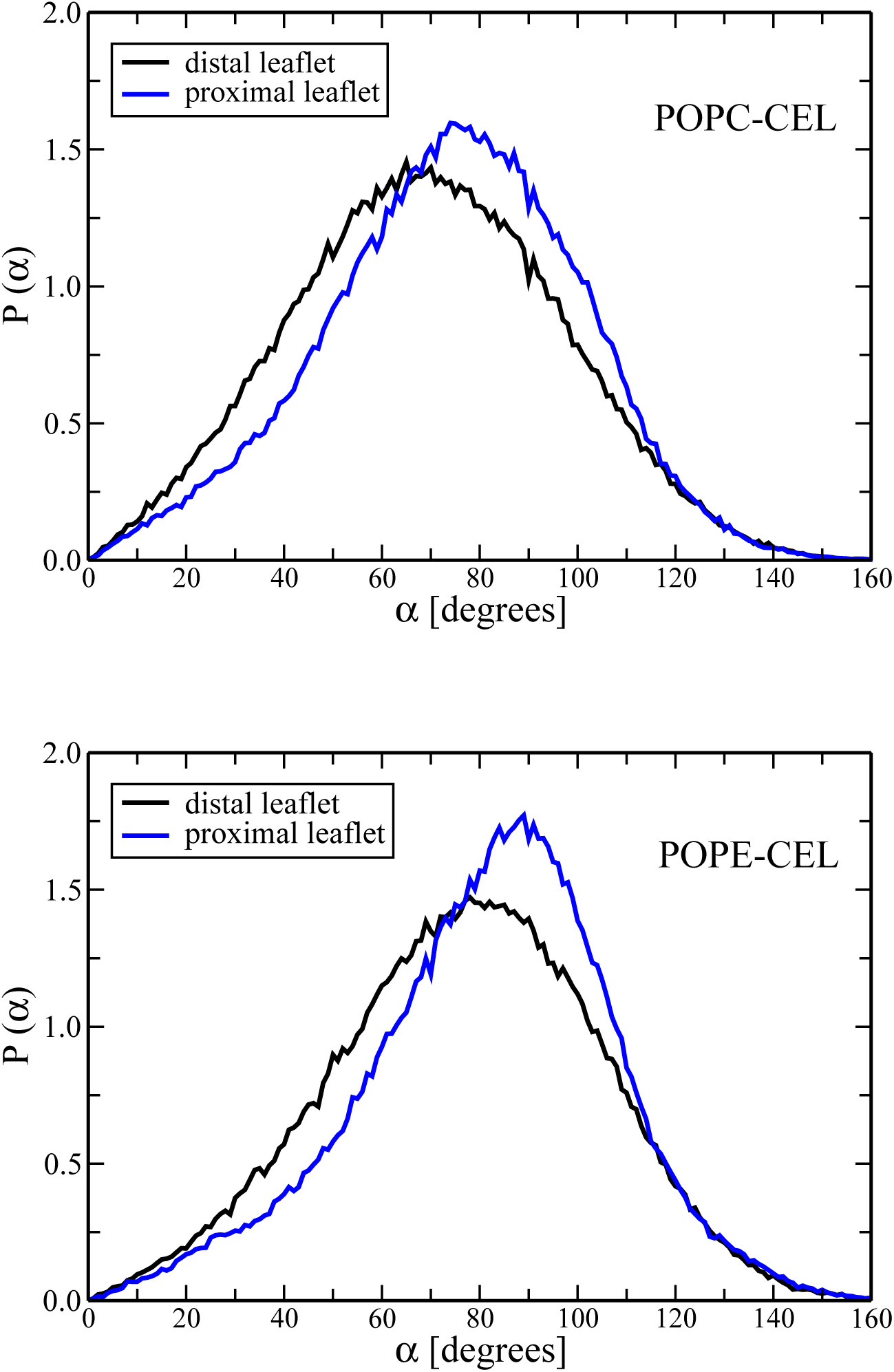
The probability distribution of the angle between PN vectors of phospholipids and the outward bilayer normal for the distal (black line) and proximal (blue lines) bilayer leaflets. Shown are results for POPC-CEL (top) and POPE-CEL (bottom) systems.

Unlike POPC lipids, polar head groups of POPE lipids are capable of forming intra-and interlipid hydrogen bonds.^45,47–49^ Since binding with cellulose results in the re-orientation of the POPE head groups, it could also affect the number of inter-POPE hydrogen bonds. Indeed, the average number of interlipid hydrogen bonds was found to be 0.81±0.01 and 0.88±0.01 bonds per POPE lipid for the distal and proximal bilayer leaflets, respectively. Therefore, cellulose promotes formation of additional hydrogen bonds within the POPE monolayer. Note that for calculating the numbers of hydrogen bonds, we used the following geometrical criteria: the donor-acceptor distance was smaller than 0.35 nm and the hydrogen-donor-acceptor angle was smaller than 30 degrees.

### Hydrogen Bonding

To gain insight into the molecular details of the phospholipid-cellulose interactions, we recall that the surface of a cellulose crystal comprise a large number of hydroxyl and hydroxymethyl groups which are both donors and acceptors of hydrogen bonds.

Table 2 summarizes the average numbers of hydrogen bonds between the phosphate groups of POPC and POPE lipids and the hydroxyl(hydroxymethyl) groups of cellulose. The number of hydrogen bonds is normalized by the total number of cellulose dimers on the crystalline surface (72 dimers).

**Table 2:**
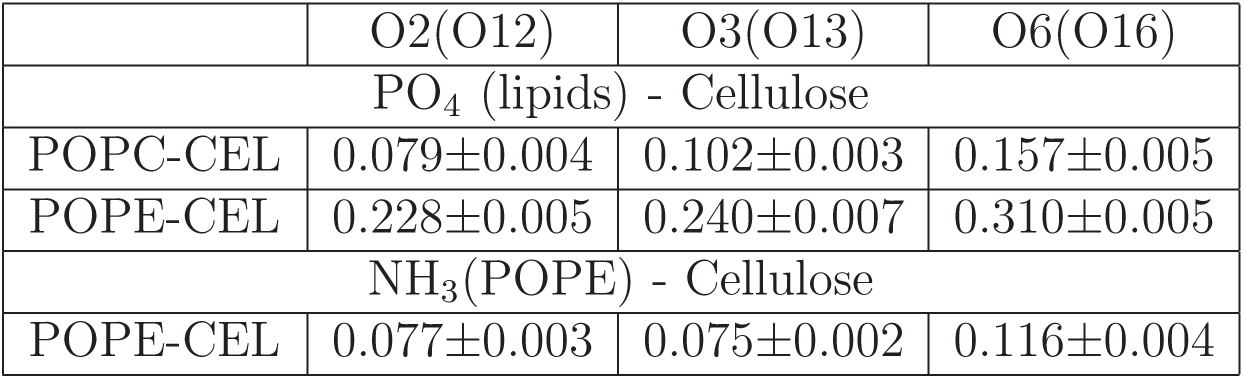
The number of hydrogen bonds between the lipid head groups (phosphate and NH_3_ groups) and hydroxyl (hydroxymethyl) groups of cellulose (per cellulose dimer).

Remarkably, the phosphate groups of POPE lipids establish considerably larger number of hydrogen bonds with cellulose than POPC lipids. In particular, our results show that the number of hydrogen bonds between POPE phosphate groups and cellulose is 2.9, 2.4, and 2 times larger for O2(O12), O3(O13), and O6(O16) cellulose groups as compared to POPC counterparts, see Table 2. Such a considerable difference can be explained by several factors: First, the NH_3_-groups of the POPE lipid head groups are smaller than the choline groups of the POPC lipids, so that they hinder access to phosphate groups to a lesser extent as compared to the POPC lipids. As a result, the cellulose hydroxyl groups can come closer to the POPE phosphate groups, Figure 5. Second, the POPE head groups are capable of hydrogen bonding with other lipids in the monolayer, so that they are oriented more horizontal with respect to the bilayer surface, which again promotes contacts between cellulose and POPE phosphate groups; this is in agreement with a previous study^50^ in which it was found that in a pure POPE membrane the P-N dipole forms an angle of about 92 degrees with respect to the normal. Third, and importantly, the NH_3_-groups of POPE lipids also can serve as donors of hydrogen bonds between POPE lipids and oxygen atoms of hydroxyl(hydroxymethyl) groups of cellulose. This type of lipid-cellulose interactions are absent in the POPC-CEL system. As Table 2 shows, the average number of the NH_3_-cellulose hydrogen bonds, being considerably smaller than that of POPE phosphate-cellulose bonds, is rather close to the number of POPC phosphate-cellulose bonds. This makes the difference in the number of cellulose-lipid hydrogen bonds for POPC-CEL and POPE-CEL even more pronounced.

If the POPE lipid bilayer forms considerably larger number of hydrogen bonds with the cellulose crystal as compared to the POPC bilayer, why is there no noticeable difference in the free-energy of binding of POPC and POPE bilayers to cellulose (see Figure 3)? To answer this question, one has to explore the influence of the cellulose binding on bilayer hydration. As discussed above, upon binding with cellulose the POPE bilayer is found to locate closer to the cellulose surface than the POPC bilayer. Therefore, one can expect lower level of hydration for the interfacial cellulose-POPE region. Indeed, our analysis shows that the average number of water molecules per lipid in the interfacial cellulose-lipid regions amounts to 6.8±0.1 and 10.6±0.1 for POPE and POPC bilayers, respectively. Therefore, binding of the POPE bilayer to cellulose is accompanied by considerably stronger dehydration, implying a larger number of broken hydrogen bonds between water molecules and lipids.

To characterize the formation and breakage of hydrogen bonds of different types, Table 3 lists the changes in the total number of hydrogen bonds which are formed/broken upon binding to cellulose (the first and last 100 ns of the 600 ns MD trajectories were considered). In line with the above findings, the number of lipid-cellulose hydrogen bonds is indeed larger for the POPE-CEL system as compared to the POPC-CEL counterpart (86 vs 24). However, upon binding with cellulose the POPE-cellulose interfacial region loses a considerably larger number of hydrogen bonds with water due to higher dehydration: 241 vs 99 broken hydrogen bonds with water for POPE-CEL and POPC-CEL systems, respectively (both lipid-water and cellulose-water hydrogen bonds are accounted). Interestingly, although water-water hydrogen bonds are not related directly to the bilayer-cellulose interactions, binding of a POPE bilayer to a cellulose crystal leads to a larger increase in the number of water-water hydrogen bonds as compared to the POPC counterpart: the more water molecules are squeezed from the interfacial region to the bulk, the more additional hydrogen bonds these water molecules can form. To further quantify the energetics of lipid-cellulose interactions, we also evaluated the changes in various components of the interaction energy (calculated as a sum of the Coulomb and Lennard-Jones energies) upon binding the bilayer to the cellulose crystal, see Table 3. In line with what was found for hydrogen bonds, the gain in the attractive lipid-cellulose interactions turned out to be twice larger for the POPE-CEL system. In turn, a positive increase in the energy due to dehydration of the interfacial region for the POPE-CEL system exceeds that for the POPC counterpart by the same factor of 2, see Table 3. Therefore, since the free-energy barriers presented in Figure 3 correspond to the differences between fully hydrated and cellulose-bound states of the lipid bilayers, an excess in the number of formed POPE-cellulose hydrogen bonds is balanced by the breakage of POPE-water and cellulose-water hydrogen bonds, leading to similar values in the free-energy of binding for both POPC and POPE lipid bilayers (−1.89 vs −1.96kJ/mol per cellulose dimer).

**Table 3:**
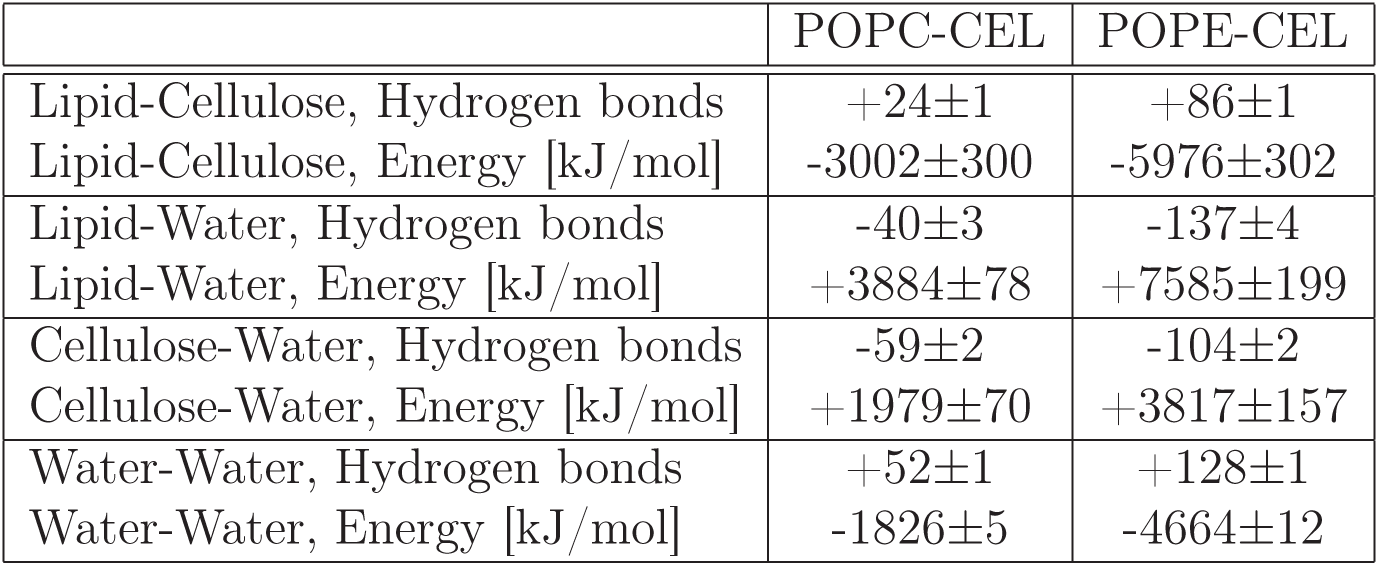
Changes in the total numbers of hydrogen bonds and in the interaction energy upon binding of the phospholipid bilayer to the surface of the cellulose crystal.

## CONCLUSIONS

The understanding of interactions of phospholipids with cellulose is of tremendous importance due to a wide use of cellulose-based materials in medicine (e.g. wound dressing and bone implants), which often implies a direct contact of a cellulose-based material with living tissue. Thus, it is critical to have detailed investigations of the impact of cellulose on cells and, more specifically, on plasma membranes that surround cells.

In this work we have performed a series of atomic-scale MD simulations of phospholipid bilayers interacting with the surface of a cellulose crystal. Lipid bilayers comprised of phosphatidylcholine and phosphatidylethanolamine lipids were studied. Both unbiased and biased umbrella sampling simulations clearly show the existence of strong attractive interactions between phospholipids and cellulose. The free energy of the cellulose-bilayer binding was found to be −1.89 and −1.96 kJ/mol per cellulose dimer for POPC and POPE bilayers, respectively. We also demonstrated that the driving force for such strong interactions is the formation of hydrogen bonds between the lipid head groups and the hydroxyl (hydroxymethyl) groups of cellulose. Interestingly, the overall number of cellulose-lipid hydrogen bonds is ~ 3.6 times larger for the POPE bilayer as compared to its POPC counterpart. However, this excess is balanced by the breakage of a larger number of POPE-water and cellulose-water hydrogen bonds, since the hydration level of the POPE-cellulose interfacial region is 1.5 times smaller than in the POPC-cellulose system (6.8 vs 10.6 water molecules per lipid). Correspondingly, the equilibrium cellulose-bilayer distance also turns out to be smaller for the POPE bilayer (0.25 nm vs 0.47 nm). Importantly, both the equilibrium hydration level of the interfacial region and the equilibrium bilayer-cellulose distance can potentially be measured in experiments.

Strong attractive phospholipid-cellulose interactions have a significant impact on the structural properties of phospholipid bilayers, resulting in asymmetry in the structures of the opposite bilayer leaflets. In particular, a cellulose crystal induces structural perturbations that are seen, e.g., in the density profiles of the phospholipids (the appearance of two peaks in the density profile instead of one and shifting lipid phosphate groups toward the crystal surface). Furthermore, we witnessed a more horizontal re-orientation of lipid head groups with respect to the bilayer surface in the lipid monolayer next to the cellulose crystal.

Such a noticeable influence of cellulose on the properties of phospholipid bilayers can be undesirable when it comes to the interactions of cellulose-based materials with cell membranes. The molecular-level insight provided by our computational study could be of great help in this respect, as major types of cellulose-membrane interactions have been identified and characterized. Our findings can be used in designing guidelines and strategies for fine-tuning the strength of the cellulose-lipid attractive interactions. In particular, the free energy of cellulose-bilayer binding could be reduced through esterification of hydroxymethyl/hydroxyl groups of cellulose,^51^ preventing thereby the formation of a large number of hydrogen bonds between the membrane and the cellulose surface. This is a subject of ongoing studies.

To conclude, it has to be emphasized that the single-component lipid bilayers considered in our study are not real cell membranes but simplified model systems implying that many important aspects of cellulose-membrane interactions are not accounted for. For instance, the strong interactions between cellulose and cell membranes are typically attributed to proteins called cellulose binding domains.^52-54^ As for phospholipid membrane-cellulose interactions, there has been surprisingly limited number of experimental studies. In particular, Setaka et al.^55^ demonstrated the possibility of the use of cellulose sheets as a versatile substrate for creating phospholipid monolayers. Furthermore, direct interactions between cellulose and phospholipids are observed in cellulose-based dialysis membranes.^56^ Given the absence of experiments that would directly address cellulose-lipid interactions, we hope that our computational findings promote such experimental studies with an aim of testing the predictions presented here. This could be achieved e.g. with modern atomic force microscopy techniques such as the recent single-lipid-extraction approach.^57^ Alternatively, one can use fluorescence and differential scanning calorimetry to probe the fluidity of phospholipid membranes.

## Supporting Information

Supporting Information is available free of charge on the ACS Publication website.

The component-wise mass density profiles for free-standing lipid bilayers; the time evolution of bilayer-cellulose distances observed in additional simulations; the chemical structure and atom numbering of a cellobiose unit.

## ACKNOWLEDGMENTS

This work was supported by the Russian Ministry of Education and Science within State Contract 14.W03.31.0014 (MegaGrant). The authors wish to acknowledge the use of the SHARCNET/Compute Canada computing facilities (Canada) and the computer cluster of the Institute of Macromolecular Compounds RAS.

